# Tissue Engineering Applications for Novel Integrated and Mobile Perfusion System

**DOI:** 10.1101/2025.03.12.642925

**Authors:** Angie Zhu, Tilak Jain, Emmett Reid, Usmaan Siddiqi, Amatullah Mir, Olivia Dunne, Narutoshi Hibino

## Abstract

Perfusion offers unique benefits to tissue-engineered systems, enhancing oxygen and nutrient transport which improves tissue formation and growth. In this study, we present a novel and integrated portable perfusion system, termed the FluidON. Weighing <10 lbs, the system can maintain continuous flow in a standard incubation environment (37°C, 5% CO_2_), effectively functioning as a portable perfusion and culture chamber. To test the system’s perfusion parameters, we measured the volumetric flow rate across a range of pressures and found that the system could achieve flow as low as 0.40± 0.19uL/s, which is similar to *in vivo* interstitial flow. Computational fluid dynamics revealed uniform flow distribution, laminar flow, and gentle circulation, helping ensure even fluid and nutrient distribution. To study the biocompatibility of the system, bioengineered tissue patches were created and perfused. Viability was assessed through flow cytometry. The system does not adversely affect cell health as the viability of perfused samples was found to be 40.99±6.22% alive after 24 hours (n=4), while that of the static control was 38.56±4.22% alive (n=4). To determine the effects of perfusion on spheroid spatial arrangement, perfused tissue patches were analyzed with light microscopy. It was discovered that perfusion promoted spheroid aggregation and cohesion, causing the distance from one spheroid to its nearest neighbor to decrease after 24 hours of perfusion. Perfusion was also found to improve the strength of hydrogels as the average hole area, caused by hydrolytic enzymes that degrade the hydrogel matrix, was smaller in perfused conditions compared to the control. Complemented by its ability to provide mobile perfusion and incubation, this novel integrated portable perfusion system holds promise for promoting tissue maturation, elevating tissue bioengineering studies.

## 1. Introduction

Tissue-engineered systems, composed of three-dimensional (3D) stem cell composites reinforced with an extracellular matrix, present promising avenues for tissue regeneration (Koda et al.). However, key concerns to clinical implementation include (1) sustained cell viability, (2) hypoxia and accumulation of toxic metabolites (Riffle et al.), and (3) development of appropriate tissue vasculature to recapitulate in vivo structures (Jarrell et al.).

Perfusion offers enhancements to these challenges; perfused conditions have been found to improve the spatial uniformity of cell distribution, enhancing the expression of specific cell markers within the core of 3D tissue aggregates (Carrier et al.). In comparison, static studies have demonstrated cell-specific markers predominantly on the periphery of tissues, reflecting the presence of nutrient concentration gradients associated with diffusional mass transport (Akins et al.). As such, perfusion facilitates increased delivery of oxygen and nutrients to the center of tissues, combating the hypoxia concerns intrinsic to the 3D nature of engineered tissues. Additionally, the fluidic movement of perfusion systems generates shear stress, creating a dynamic mechanical environment that enhances the generation of vasculature and tissue development (Strobel et al.).

Perfusion systems can generally be classified as microfluidic, interstitial, or organ, with each targeting a specific construct. Microfluidic perfusion systems target single organoids with sub-microliter volume flow rates (McLennan et al.). Interstitial systems aim to mimic biological interstitial flow rates in tissues. Finally, large organ perfusion generally tailors to larger models, such as whole organs (Macdonald et al.). However, across these systems, similar limitations arise. Because perfusion requires the usage of pump systems—which are often large and heavy—perfusion systems generally lack mobility which makes transportation of engineered tissue and broader applications past the laboratory difficult. Moreover, laboratories often must design their own perfusion systems, using a variety of external tools. While this allows researchers to tailor their systems to their unique needs, this also limits the reproducibility of studies across different facilities. A portable and easy-to-use perfusion system is still missing. As such, we hereby detail the use of a novel portable and modular perfusion system to obtain precise pressure control to achieve media flow **(Fig. 1A)**.

**Figure 1:**
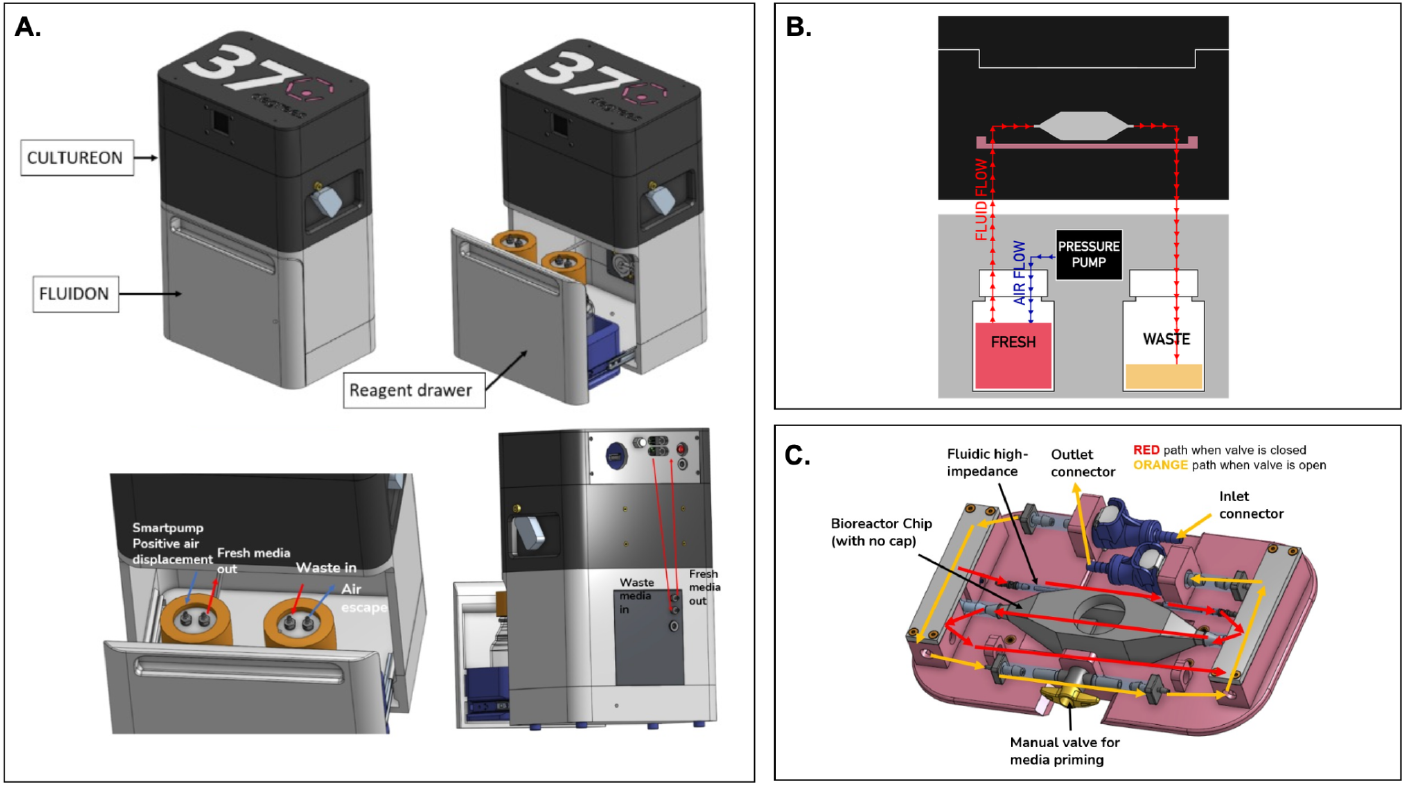
An overview of the integrated portable perfusion system. **(A)** As shown by a computer-aided design (CAD) of the perfusion system, the CultureON and the FluidON are integrated. A reagent draw in the FluidON hosts the fresh and waste bottles while a Smartpump allows for continuous media flow. **(B)** A simplified perfusion system schematic demonstrates how pressure generated in the FluidON routes media flow into the CultureON, where the bioreactor chip houses the tissue. (C) Represented in a CAD, a custom bioreactor tray and chip were designed. Depending on the valve orientation, two flow paths are possible. One bypasses the bioreactor chip, allowing for quick fluid line priming. The other route the fluid through a tubing constrictor, tuning the flow to create low biomimetic flow rates. This low flow is then passed through the bioreactor chip.

## 2. Results

### a. Design of FluidON and Perfusion System Components

#### FluidON and CultureON

This novel perfusion system—called the FluidON—is pressure-based. A general pressure-based perfusion system connects a perfusion chip in series with two reservoirs. Reservoir 1 begins perfusion filled with media while Reservoir 2 is empty. Perfusion occurs as air enters Reservoir 1 and applies pressure on the surface of the media in the bottle. This causes the media to flow out of the reservoir through silicon tubing and the perfusion bioreactor where it finally deposits in Reservoir 2.

The FluidON (37degrees, Inc.) likewise utilizes airflow for perfusion **(Fig. 1B)**, but several important modifications allow for precise control of fluidic flow. The FluidON module utilizes a piezoelectric air pump that drives filtered air (1.2 microns) into a fresh reagent bottle. A 100-micron bore (I.D) 4-inch tubing constrictor tunes the media flow, providing high-fluidic impedance and allowing for desired low flow rates. Media continuously flows throughout the FluidON system, into the bioreactor, and is ultimately collected via an outlet to the waste container. Furthermore, flow is routed to the CultureON (37degrees, Inc.)—a portable CO_2_ incubator module—where the bioreactor is housed in a bioreactor tray assembly. The CultureON module controls the temperature at 37°C and 5% CO_2_, maintaining the bioreactor components at standard culture conditions.

The dual integration of FluidON and CultureON simultaneously provides optimized flow and portable culture conditions. The CultureON module uses disposable CO_2_ gas cartridges and is DC battery-powered, removing the need for external gas supply lines. Additionally, the CultureON module is designed to stack vertically on the FluidON module, which contains a reagent drawer that houses the fresh media and waste bottles. The CultureON unit can be completely disconnected from FluidON so that the bioreactor can be transported without disrupting the culture conditions. The total dimensions of the system fit within 14 x 9 x 6 inches, taking up a minimal bench footprint. Both modules weigh < 5 lbs individually, facilitating easy transport through laboratory and clinical settings.

#### Bioreactor Tray and Chip

A custom-designed tray (37degrees, Inc.) houses the bioreactor **(Fig. 1C)**. Once the inlet and outlet connections are made, the manual valve can be opened or closed. When the valve is open, media flow bypasses the high-impedance fluidic resistor and the bioreactor for fast fluid line priming of the entire system. This fills the fluid lines and ensures bubbles are removed from the system. When the manual valve is closed, the media flows through the high-fluidic impedance, which significantly reduces the flow rate as desired, and passes media through the bioreactor. The setup can maintain flow in the CultureON incubator for several days, with the volume of fluid initially added acting as the limiting reagent.

The bioreactor is composed of two components: the main chamber and the lid. Before connection to the FluidON, the tissue sample can be loaded into the chamber, and the lid is secured with parafilm to create an airtight seal that holds the two components together **(Fig. 2A,B)**. The main chamber can hold a 10 mm x 10 mm x 10 mm (1 cm^3^) scaffold in the center compartment. The inner compartment is fitted with posts that hold the patch in place. The two walls of the inner compartment that face the inlet and outlet ports are porous, such that fluid can easily flow through the patch without shifting its position.

**Figure 2:**
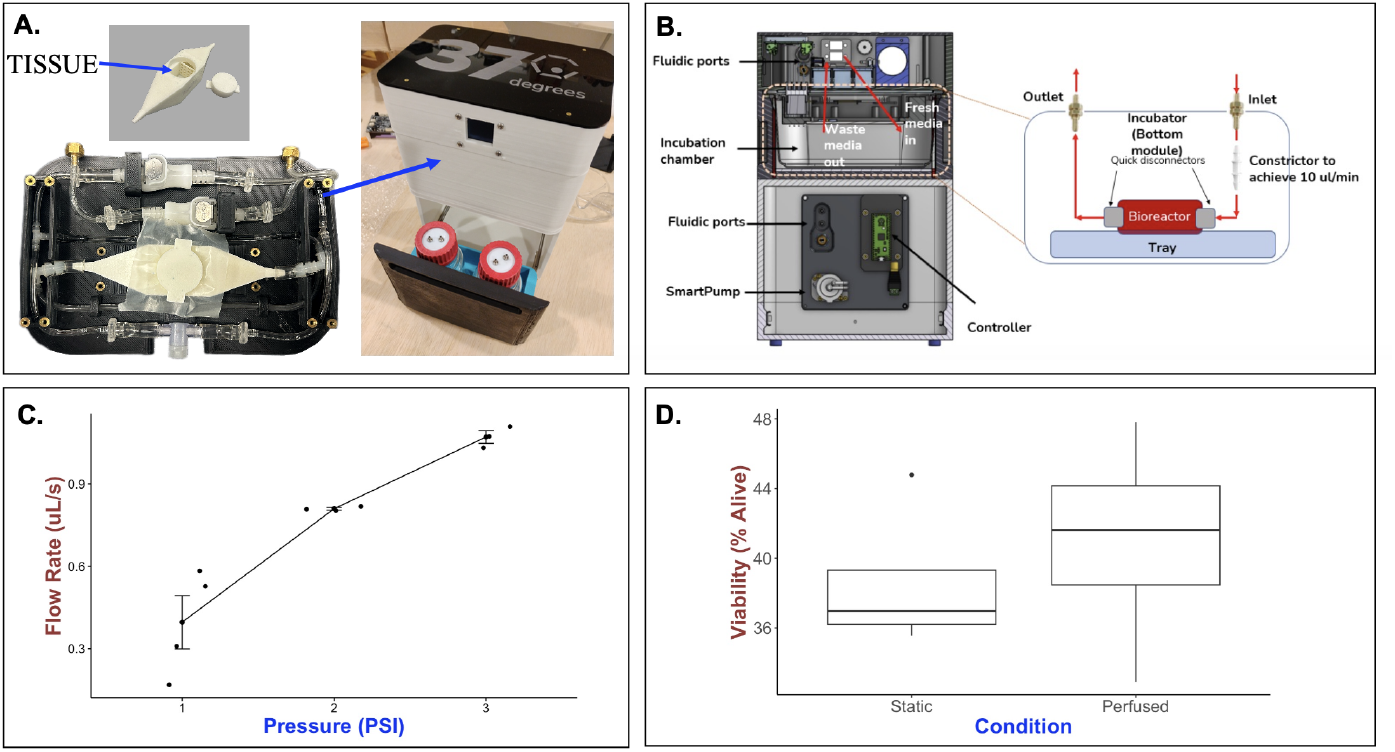
Real-time performance of the integrated perfusion system. (A) The integrated perfusion system (right) and custom bioreactor try and chip (left) takes up minimal lab bench space. **(B)** A detailed CAD shows the internal workings of the system. To test the mechanical parameters of the FluidON, **(C)** volumetric flow rate was determined across a range of pressures. It was found that the system encompasses flow as low as 0.40± 0.19uL/s. To determine the compatibility of the system with live cells **(D)**, 33k mMSC and mEC spheroids were formed and suspended in a mixture of thrombin and fibrin to form a patch. This patch was perfused for 24 hours, and it was found that the system did not adversely affect cell viability. The average viability after 24 hours was 40.99±6.22% alive while that of the static control was 38.56±4.22% alive (p-value = 0.55).

The current model of the FluidON contains media bottles that can be disconnected for sterilization through high-temperature autoclave. We perfused 70% ethanol throughout the system for 15 minutes to sterilize other remaining non-autoclavable components. After 24 hours of perfusion with culture conditions maintained in the CultureON, we observed no contamination.

### b. Experiments to Determine System Properties and Tissue Compatibility

#### FluidON Perfusion Parameters

To determine the FluidON’s flow rate as a function pressure, we measured the volumetric flow rate (V) across a range of pressures **(Fig. 2C)**. We found that the system could achieve flow rates as low as 0.40± 0.19uL/s at 1 PSI. Equation 1 was used to calculate the average flow velocity through the tissue sample containing portion of the bioreactor at 1, 2, and 3 PSI (Fig. 3) Applying this minimum flow rate to equation 1 where A_c_ is the cross-sectional area of the bioreactor (234mm^2^) yields a minimum average flow velocity (u) of 1.7±0.83µm/s.

**Figure 3:**
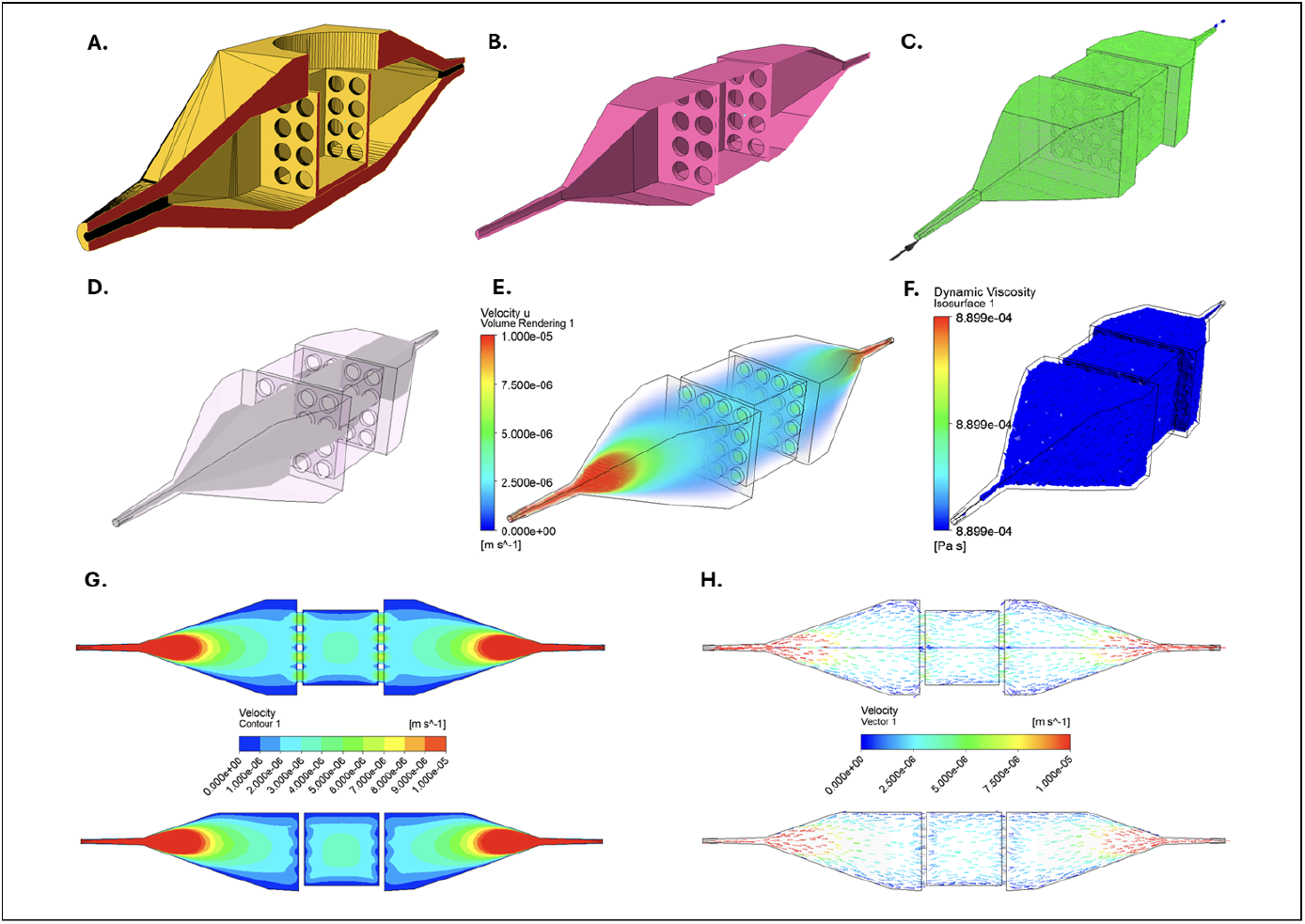
Overview of bioreactor computational fluid dynamics. (A) Modeled geometry of the perfusion chamber. (B) Visualization of the flow domain surfaces, showing where the fluid contacts the inner walls of the device. (C) Computational mesh and boundary condition setup in ANSYS CFX, after extraction of the flow domain, with refinement around the orifices and inlet/outlet transitions. (D) Extraction of the flow domain (internal volume), additionally illustrating two axial reference planes for subsequent analysis of flow patterns. (E) Velocity volume-contour plot illustrating both the acceleration and deceleration of flow (red plumes). (F) Dynamic viscosity iso-surface plot showing viscous-flow-dominated regime. (G) Two cross-sectional orientations of velocity contours on the reference planes demonstrate how the flow accelerates and decelerates in the respective regions. (H) Cross-sectional velocity vectors confirm uniform flow distribution, laminar behavior, and gentle circulation within the specimen chamber. Analysis was performed in ANSYS CFX under laminar conditions, assuming a Newtonian fluid equivalent to the culture media.

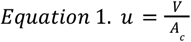

This is 2.8-5 times the average interstitial flow velocity of native muscle tissue of 0.34-0.6µm/s (Chary et al.) (Fukui et al.). Furthermore, we can use equation 2 to calculate the Reynolds number given u in cm/s (1.7×10^-8^ cm/s) the media density (? = 1.009±0.003g/cm^3^), media dynamic viscosity (µ = 0.930±0.034mPa·s) and the hydraulic diameter of the bioreactor (D=1.5cm) (Poon).

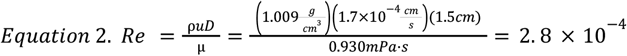

With a Reynolds number of 2.8×10^-4^, the flow through the tissue-containing compartment of the bioreactor is laminar (2.8×10^-4^ < 2300), and creeping flow conditions apply to the Navier-Stokes equation (2.8×10^-4^ < 1).

Computational fluid dynamics (CFD) analysis demonstrates, visually, the flow patterns within the bioreactor **(Fig. 3)**. Fluid velocity is shown to be evenly distributed in the chamber that holds the tissue sample. Dynamic viscosity analysis shows that the isosurface encompasses the entire internal volume, indicating that the fluid fully occupies all passages without separation or stagnation. This validates a viscous flow-dominated regime suitable for cell perfusion. Likewise, cross-sectional velocity vectors confirm uniform flow distribution, laminar behavior, and gentle circulation within the tissue holding chamber, which benefits even nutrient exposure. The FluidON, therefore, can be categorized as a perfusion system targeting interstitial flow rates. Balanced fluid distribution, velocity, and viscosity within the tissue chamber ensure uniform perfusion of samples, supporting consistent nutrient delivery and helping promote tissue maturation.

#### Effects of Perfusion on Bioengineered Tissues

Effects of the FluidON system on bioengineered tissue samples were also determined. Hydrogel patches were created by embedding mouse mesenchymal stem cells (mMSC) and mouse endothelial cells (mEC) spheroids in fibrin and thrombin. The mixture then underwent gelation, forming a semi-solid structure. This patch was subject to perfusion in the FluidON.

Patch viability was measured using flow cytometry. The viability of samples perfused for 24 hours was found to be 40.99±6.22% alive (n=4). In comparison, the static samples were 38.56±4.22% alive (n=4) **(Fig. 2D)**. A t-test returned a p-value of 0.5455 (t = 0.64, df = 5.27). The FluidON system successfully achieves low flow rates without adversely affecting cell viability.

Light microscopy was also used to assess the effects of perfusion on the patch as perfusion presented benefits to tissue quality **(Fig. 4A)**. The area of matrix metalloproteinase induced holes in the hydrogel was quantified after 24 hours in perfused (n=3) and static conditions (n=3) **(Fig. 4B)**. It was observed that the hole area decreased in perfused conditions compared with static conditions. Perfusion resulted in holes of an average pixel area of 5.51e-2±1.28e-3 per spheroid while that for static conditions was 7.03e-3±2.48e-4 (p-value = 0.17).

**Figure 4:**
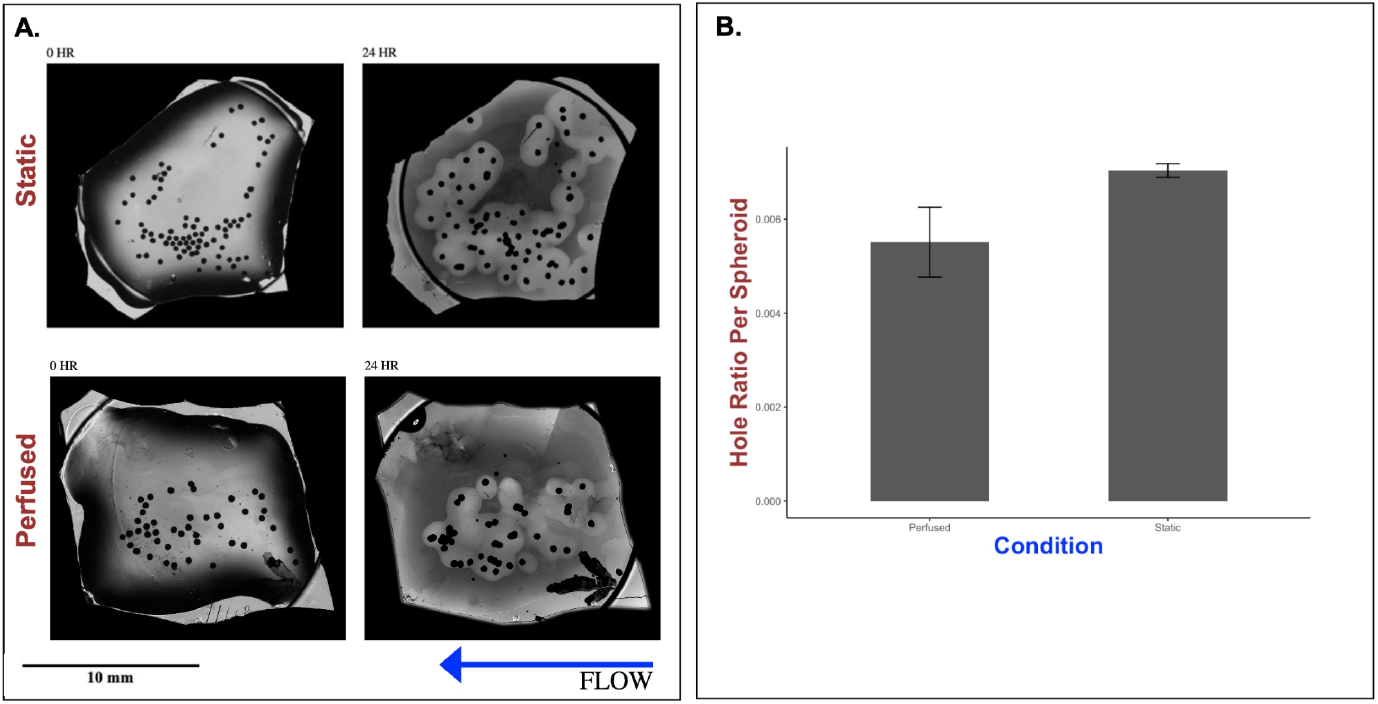
Effects of perfusion on hydrogel strength. To visually inspect the effects of perfusion on engineered tissue, 33k mMSC and mEC spheroids were formed and suspended in a fibrin and thrombin mixture. This mixture was plated on a glass coverslip and perfused for 24 hours. **(A)** Light microscopy was used to analyze the effects of perfusion on the patch. Holes were observed to form in the hydrogel matrix after 24 hours. **(B)** Quantitative analysis of the hole area demonstrated that perfusion reduced hole size to an average pixel area of 5.51e-2±1.28e-3 (n=3) per spheroid compared to 7.03e-3±2.48e-4 per spheroid (n=3) observed in the static control (p-value = 0.17).

Spatial changes were analyzed to see if perfusion affected the arrangement of spheroids. After setting spheroids in the hydrogel, it was observed that 24 hours of perfusion promoted cohesion and clumping compared to the static control where spheroids were more widely dispersed. Analysis using Fiji showed that distance to the nearest neighbor decreased from 0.60±0.22mm to 0.54±0.34mm after 24 hours with perfusion, while that of the static condition grew from 0.55±0.19mm to 0.75±0.43mm **(Fig. 5A)**. Natural variability arises during the creation of tissue patches because the methodology used to suspend spheroids in the hydrogel is inherently random. To discount outcomes resulting from chance, the data was normalized to the average distance at the 0-hour time point **(Fig. 5B)**. After normalization, the trend of increased cohesive clumping remained in the perfused condition; while the normalized distance grew in the static control, it decreased in the perfused sample.

**Figure 5:**
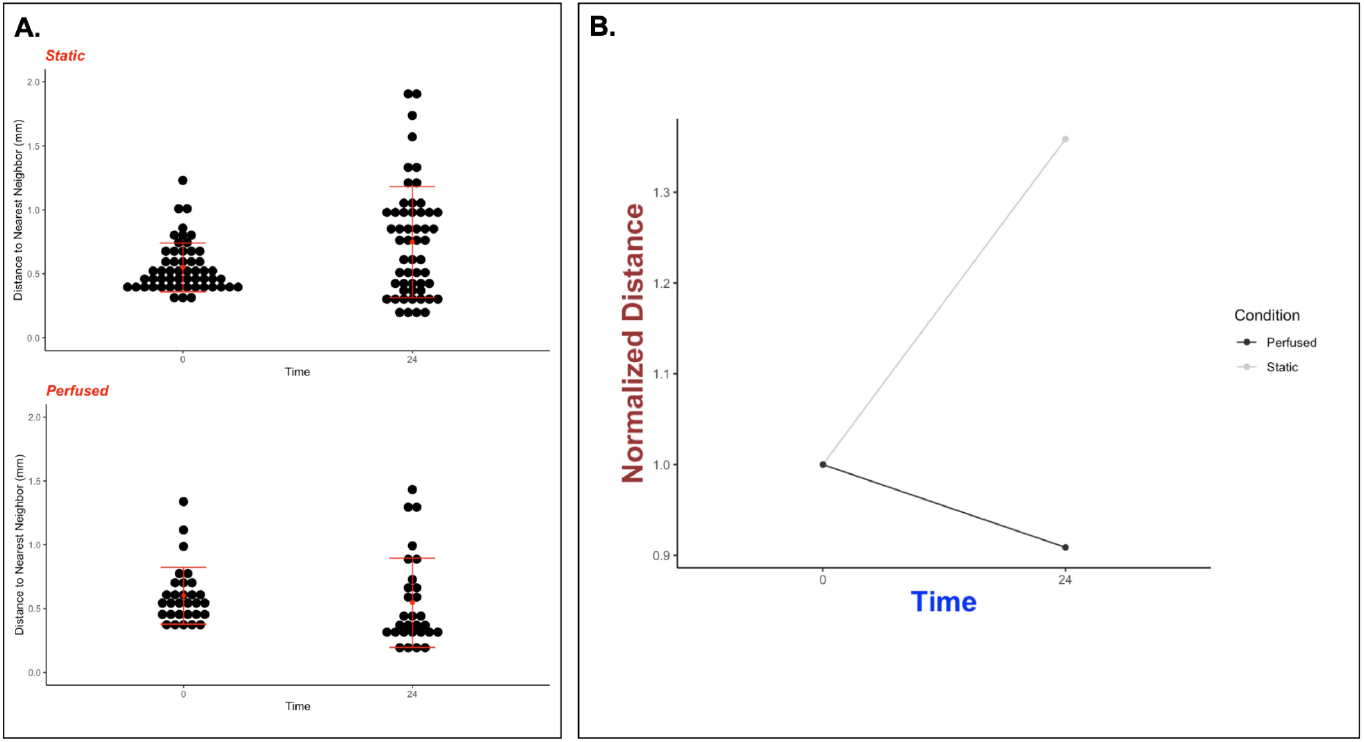
Effects of perfusion on spheroid spacing. To determine spatial changes caused by perfusion, tissue patches were analyzed with and without perfusion. **(A)** Distance from each spheroid to its closest neighbor spheroid was measured for all spheroids in the sample. It was found that after 24 hours, the spheroids in the perfused sample remained more cohesively clumped relative to the static sample where spheroids were more widely dispersed. **(B)** After normalizing the data to account for chance, the same trend of increased cohesion and clumping in perfused samples remained as the distance from a spheroid to its nearest neighbor decreased.

## 3. Materials and Methods

### Cell Culture

Mouse mesenchymal stem cells (mMSC) and mouse endothelial cells (mEC, C166-GFP, ATCC) were cultured in media constituted of Dulbecco’s modified Eagle’s medium (DMEM), 10% fetal bovine serum (FBS), and 1% penicillin/streptomycin. Cultures were incubated at 37°C and 5% CO2 for 3-4 days.

### Spheroid Creation and Dissociation

To create spheroids, 33,000 total cells––at a ratio of 80:20 mEC to mMSC–– were added to each well of a 96-well (U) bottom plate (IWAKI) along with 200μL of media. The plate was incubated for 72 hours at 37°C and 5% CO_2_. Spheroids were harvested and transferred to a 6-well plate for dissociation. 5mL collagenase (255 U/mg dry weight, 2 mg/mL stock concentration) was added, and the plate was incubated for 24 hours. Post-incubation, spheroids were pipetted up and down to introduce shear force.

### Hydrogel creation

Thrombin from human plasma and fibrinogen from human plasma (Sigma) were diluted to 20 Units/mL and 40 mg/mL aliquots, respectively. To create a patch for microscopy, roughly 100 spheroids were suspended in 75uL of diluted thrombin before being combined with 75uL of diluted fibrin. This mixture was pipetted onto a glass coverslip modified to roughly fit 1cm X 1cm dimensions, where surface tension maintained its 3D structure. Variability in the number of spheroids suspended may occur as spheroids adhere to the pipette tip, resulting in partial transfer.

To create a patch for flow cytometry analysis, 300 spheroids were suspended in 500uL diluted thrombin before combination with 500uL of diluted fibrin. This mixture was pipetted into a disposable 10x10x5 mm vinyl specimen mold (Sakura). Once hydrogel was set, the vinyl mold was cut and the patch was carefully removed with an X-ACTO knife. For static conditions, the patch was incubated in a 35mm glass bottom dish (Cellvis) with 3mL media. For perfusion, the patch was inserted into the FluidON bioreactor chip and subject to perfusion at 2 PSI.

### Flow cytometry

Spheroids post-dissociation were transferred to 5mL 12x75mm tubes and washed to remove the supernatant. Cells were further washed and resuspended in 1mL Phosphate-buffered saline. To stain for dead cells, 1uL LIVE/DEAD™ Fixable Blue Dead Cell Stain Kit (Invitrogen™) was added to each tube. Samples were then washed to remove excess stain and fixed with 1000μL 10% neutral buffered formalin for 15 minutes. Fixed cells were then washed in 1mL of 1% Bovine serum albumin. In an ice box, cells were brought to the Cytometry and Antibody Technology Core Facility (University of Chicago) where the NovoCyte Pentron Flow Cytometer was used.

### Analysis

Patches were analyzed with light microscopy on an Invitrogen EVOS microscope. Images were processed in Fiji. Hole size was found by measuring the pixel area of each hole. This value was divided by the number of spheroids in the initial patch to account for variability in the spheroid plating number.

The distance to the nearest neighbor was defined as the distance from the center of one spheroid to the center of the closest spheroid. To normalize the data, the average distance to the nearest neighbor at 0 hours was calculated and scaled to a value of ‘1.’ The distance at 24 hours was divided by the average at 0 hours, and this new ratio was compared to the scaled value of ‘1.’ If the ratio was >1 it was determined that the distance to the nearest neighbor grew, and if the ratio was <1 it was interpreted that the distance to the nearest neighbor decreased.

R Studio was used for numerical and graphical analysis. A two-tailed Student’s t-test was performed for statistical comparison.

### Computational Flow Dynamics Analysis

CFD simulations were conducted in ANSYS CFX to model flow through the perfusion chamber. The fluid domain was extracted from the chamber geometry, and a tetrahedral mesh was generated with refinement around the inlet, outlet, and orifice regions. The fluid was modeled as incompressible and Newtonian, with properties approximating standard cell culture media at 37 °C (density and viscosity values based on literature or prior measurements). A steady-state, laminar flow regime was assumed. The inlet boundary condition was set as a mass flow rate of 4.0 × 10^−7^ kg/s, reflecting typical operating conditions for the chamber, with static pressure of 1 atmosphere set at the outlet. Solutions were considered converged when residuals dropped below 1 × 10^−6^, and flow characteristics were evaluated through velocity contours, vector fields, and viscosity iso-surfaces to confirm full perfusion without separation.

## 4. Discussion

Perfusion presents a novel means to enhance bioengineered tissues, and many unique systems have been introduced in recent years. One such is the organ-on-a-chip (OoC) model, which allows for biomimetic flow and the integration of key mechanical and electrical stimuli. However, because the system is constituted by microfluidic devices, it is tailored toward single organoids and microtissues rather than tissue constructs (Cruz-Moreira et al.). On the other hand, large organ perfusion systems house larger tissues or organs, but their greater flow velocity may harm tissue maturation (Calder et al.). Perfusion systems that target interstitial flow combine features of both Organ-on-a-Chip and large organ perfusion; they provide slow, biomimetic flow while offering space to support tissue constructs. Therefore, while OoC and large organ perfusion have their unique uses, interstitial flow provides an ideal niche for tissue growth and maturation.

The FluidON, introduced in this paper, targets medium interstitial flow rates to support 3D tissue formation. A study by Radisic et al. similarly details a perfusion system with interstitial flow. They were able to perfuse a tissue scaffold for 7 days at 0.5ml/min, and they noted significant increases in viability following perfusion relative to their control (Radisic et al.). A slower flow rate of 0.045ml/min was used in this study, but the FluidON can be tailored to the desired flow rate by changing the pump pressure. Likewise, though our experiment was only conducted for 24 hours, the desired test length can be increased as long as there is sufficient media. Radisic et al. noted positive effects of interstitial perfusion on cell viability. Though the FluidOn demonstrated no significant changes in cell viability after 24 hours, with longer study times, we anticipate similar improvements in viability, reflecting the system’s potential to support cell survival and tissue growth.

Regardless of price point or flow targeted, most established perfusion systems use a variety of pumps, but they are typically external or large (Sierad et al.) (Grayson et al.) (Bender et al.). Additionally, long-term perfusion experiments also require co-usage with an incubator, further increasing the size burden of the perfusion system. Taken collectively, this limits the portability of perfusion systems, anchoring them to a laboratory bench and reducing broader applications.

There lacks a system able to maintain perfusion under culture conditions for long durations, which could be useful for surgical specimen collection, transportation of samples from one laboratory to another, and more. Our perfusion system may provide a potential remedy as it integrates perfusion and culturing in a construct weighing <10 lbs. In their study, Wolf et al. introduces a portable perfusion system—the VascuTrainer—which is able to achieve pulsatile flow ranging from 10 to 2000 mL/min with a battery life of approximately 25 hours (Wolf et al.). Though demonstrating positive results, the VascuTrainer operates at a much higher flow rate than the optimal for interstitial flow as well as the FluidON. Additionally, it lacks the ability to support tissue culture during transport, making it better suited to receiving organ implants than tissue maturation. Able to perfuse and culture simultaneously in a compact and mobile system, the FluidON may target portable tissue maintenance better than other available counterparts.

While preliminary studies demonstrate FluidON’s promising application, as an initial prototype, limitations may arise. To begin, though our results show meaningful differences, they are not statistically significant. We believe this is due to the short duration of our experiments, and longer studies could demonstrate more robust results. In addition, as the current FluidON version did not have built-in refrigeration, the fresh media bottle had to be maintained in an external ice container for the duration of the experiment. The media, however, warms up to 37°C as it is routed into the CultureON incubation environment and before entering the Bioreactor. Other perfusion systems place the fresh media source in an incubator alongside the sample. However, prolonged exposure to warm conditions can degrade metabolites in the culture media, reducing its effectiveness. Integration of refrigeration into FluidON reagent bay will allow for an easier experimental setup and allow for increased perfusion durations (days to weeks).

Additionally, while the system pressure and drive voltage can be easily examined through an external monitor, new additions can be added to further improve the capabilities of the system. Precision flow sensors situated at crucial junctions in the system could enable real-time monitoring of the internal fluidic performance of the perfusion circuit. Moreover, development of a live video microscopy system is underway, which could potentially reveal new insights into bioengineered tissues. Although most perfusion studies run continuously over time, analysis is typically limited to discrete time points, rather than capturing real-time data. Integrating a live video microscopy system into the CultureON and FluidON would help us understand cell migration patterns and more in real-time to better understand how perfusion impacts bioengineered tissues.

## 5. Conclusion

In this study, we introduced a novel and integrated portable perfusion system. Composed of the CultureON, the FluidON, a custom-designed bioreactor tray, and a bioreactor chip, the system is capable of maintaining continuous fluid flow at 37°C and 5% CO_2_. While the incubation environment is similar to that of standard incubators, a significant difference is that the total weight of the new solution is < 10 lbs. The components of the system stack on each other, and the FluidON is designed with fresh and waste media storage compartments, facilitating easy and secure transportation between locations. Using a high-impedance fluidic resistor, the system achieves low flow velocity—similar to that of interstitial flow. However, its fluidic performance allows for a 1 cm^3^ scaffold, expanding its utility from previously reported organ-on-a-chip and single-spheroid perfusion systems.

This system is a new perfusion method and integrated design suitable for a variety of bioengineering studies. Preliminary studies have demonstrated the system’s promise in elevating tissue engineering outcomes; viability was not affected while the spatial arrangement and properties of the hydrogel favored the development of enhanced tissue patches. While the current applications of the FluidON are targeted toward bioengineered tissue perfusion, further applications can be envisioned. Given its portable nature and its ability to sustain culture conditions with minimal bench space, the FluidON could be potentially extended toward improving specimen collection, research transportation, and organ transplant health. While further studies must be conducted, this novel perfusion system shows promise for tissue engineering applications and more.

## Supporting information

Original Data for Figures

## 6. Acknowledgments

The authors would like to gratefully acknowledge Dr. Tong-Chuan He for his generous support in providing our mouse mesenchymal stem cells. The authors would also like to extend gratitude to William Stein for his diligence in experimental assistance and to Noor Syafira Shih for her support in designing the custom bioreactor.

## 8. Author Contributions

Conceptualization, N.H., T.J.; Methodology, A.Z.; Software, T.J. and U.S.; Validation, A.Z., E.R.; Formal analysis, A.Z., E.R., A.M.; Investigation, A.Z.; Resources, N.H., T.J.; Data curation, A.Z., E.R.; Writing---original draft preparation, A.Z., O.D.; Writing---review and editing, A.Z., E.R, T.J, N.B; Visualization, A.M., A.Z., U.S.; Supervision, N.H.; Project administration, N.H., T.J; Funding acquisition, N.H. All authors have read and agreed to the published version of the manuscript.

